# A Comprehensive Benchmark of EEG-Based BCI Deep Learning Models for MCI and Dementia Classification

**DOI:** 10.64898/2026.08.12.743255

**Authors:** Vasilii Zaitsev, Chun-Shu Wei

## Abstract

Electroencephalography (EEG) is a promising tool for automated detection of mild cognitive impairment (MCI) and dementia, but comparisons across studies are limited by inconsistent datasets and evaluation protocols. This study benchmarks ten deep learning models across four resting-state EEG datasets and eight binary classification tasks using a unified preprocessing pipeline and five-fold subject-wise cross-validation. Each experiment was repeated ten times. SCCNet obtained the highest mean subject-level accuracy, sensitivity, and F1 score, while ShallowConvNet achieved the highest mean segment-level accuracy, specificity, and precision. Subject-level aggregation improved mean accuracy for all evaluated models, and performance varied substantially across datasets and diagnostic tasks. Higher computational cost did not consistently correspond to better classification performance, with several compact architectures remaining competitive with substantially larger models. The results provide a reproducible reference for comparing EEG-based dementia classification models under consistent subject-independent evaluation conditions.

## 1 Introduction

Early cognitive decline may be difficult to identify because measurable cognitive changes may appear before a person loses independence in daily life (Gauthier et al., 2006). Mild Cognitive Impairment, or MCI, refers to this stage, in which cognitive performance is lower than expected for normal aging but does not yet reach the level of cognitive disability (Gauthier et al., 2006; Mosti et al., 2019). MCI is clinically important because affected individuals have a higher risk of later developing dementia, especially Alzheimer’s disease (Roberts et al., 2014; Livingston et al., 2020). However, diagnosing MCI remains challenging because it is largely dependent on neuropsychological tests, functional evaluation, clinical history, and expert judgment (American Psychiatric Association, 2013; Albert et al., 2011). These limitations highlight the need for objective computational biomarkers that can support more reliable and reproducible early detection (Jack et al., 2018).

In current clinical practice, cognitive impairment is usually assessed through cognitive tests, functional evaluations, and medical history (American Psychiatric Association, 2013; Alzheimer’s Association, 2024). When further evidence is needed, clinicians may use biological and neuroimaging markers, such as cerebrospinal fluid amyloid-beta and tau proteins, as well as positron emission tomography and functional magnetic resonance imaging (Symms et al., 2004; Jack et al., 2018; Yin et al., 2013). These methods can provide important information about pathological and functional changes in the brain (Scheltens et al., 2021). Nevertheless, they are often difficult to apply in routine screening because they can be invasive, expensive, time-consuming, or dependent on specialized clinical facilities (Zeng et al., 2024). This limits their usefulness for repeated monitoring and large-scale assessment.

Electroencephalography (EEG) provides a complementary approach for investigating cognitive decline because it measures brain electrical activity in a non-invasive, relatively inexpensive, and accessible way (Sanei and Chambers, 2013). Resting-state EEG is particularly useful for MCI and dementia research, as changes in spontaneous neural activity may reflect early functional disturbances related to neurodegeneration (Gaubert et al., 2019; Babiloni et al., 2021; Yang et al., 2019). These alterations may appear before clear structural brain atrophy or severe cognitive symptoms become visible (Cassani et al., 2018). Together with recent advances in machine learning and deep learning, EEG-based analysis has therefore become a promising direction for automated detection of MCI and dementia (Aviles et al., 2024; Azami et al., 2025).

However, the real-world implementation of the automated EEG-based BCI for MCI and dementia diagnosis using the existing methods remains limited by several methodological issues (Wang et al., 2026). The major concern is the evaluation protocol. For real-world clinical deployment, the model’s generalization to unseen subjects is necessary, which requires applying subject-level evaluation for a more realistic assessment of the diagnostic performance (Kunjan et al., 2021; Lee et al., 2023). However, in most studies that propose EEG-based MCI/dementia diagnosis methods, the EEG recordings are commonly divided into multiple epochs, and only the segment-wise splitting is done, which might allow data from the same subject to appear in both training and testing datasets (Brookshire et al., 2024). This may lead to subject-specific information leakage and inflate reported performance to the suspiciously nearly-perfect level (Young et al., 2025). Another limitation is that many studies report results from a single experimental setting, often without repeated runs or systematic comparison against strong EEG classification baselines (Miltiadous et al., 2026). This makes it difficult to assess whether reported performance is stable across random initialization, fold assignment, and model selection (Wang et al., 2026; Miltiadous et al., 2026). Moreover, EEG dementia datasets differ in cohort size, diagnostic labels, recording duration, sampling rate, and channel configuration (Cassani et al., 2018; Perez-Valero et al., 2021). Applying the same evaluation protocol to multiple datasets can therefore provide a broader and fairer view of model behavior, even when each dataset is evaluated independently (Wang et al., 2026).

To address these issues, this study provides a reproducible within-dataset, subject-independent benchmark of ten EEG deep learning models across four resting-state EEG dementia datasets. All models are evaluated using a unified preprocessing, training, and cross-validation protocol, while ensuring that subjects are kept separate across training, validation, and testing folds. Both segment-level and subject-level performances are reported, and the experiments are repeated multiple times to assess the stability of the results.

The main contributions of this study are: (1) a unified evaluation of ten representative EEG deep learning architectures across four MCI- and dementia-related EEG datasets and eight binary classification tasks; (2) a subject-independent evaluation protocol that prevents EEG segments from the same subject from appearing across training, validation, and test partitions; (3) a comparison of segment-level and aggregated subject-level performance under repeated experiments; (4) a comparison of computational efficiency in terms of trainable parameters, training and inference time, and GPU memory usage; and (5) an openly available implementation that facilitates reproducible comparison of deep learning models for EEG-based MCI and dementia classification.

## 2 Methods

### 2.1 Datasets

To evaluate the DL models across heterogeneous dementia-related EEG data, four datasets containing resting-state EEG recordings were included in this study: GENEEG Mishra et al. (2025), MCIHC Kashefpoor et al. (2019), ADFTD Miltiadous et al. (2023), and CAUEEG Kim et al. (2023). Their principal characteristics are summarized in Table 1. GENEEG and MCIHC datasets were both used for MCI vs HC classification tasks, with two available MCIHC cohorts combined into one dataset to increase the amount of data available for the models’ benchmarking. For the ADFTD dataset, three binary classification tasks were constructed: AD vs HC, FTD vs HC, FTD vs AD. Similarly, Dementia vs Normal, MCI vs Normal, and MCI vs Dementia were constructed from the CAUEEG dataset.

**Table 1:** Summary of the evaluated EEG datasets. MCI – Mild Cognitive Impairment; HC/Normal – Healthy Controls; AD – Alzheimer’s Disease; FTD – Frontotemporal Dementia.

| Dataset | Group | Subjects | Age (years) | Channels | Original Sampling Rate (Hz) | Mean Session Duration (min) |
| --- | --- | --- | --- | --- | --- | --- |
| GENEEG | MCI | 28 | $73.4 \pm 9.9$ | 17 | 200 | 4 |
| | HC | 35 | $73.4 \pm 4.7$ | | | |
| MCIHC | MCI | 16 + 16 | $68.5 \pm 8.5$ | 19 | 256 | 31.08 |
|  | HC | 11 + 18 |  |  |  |  |
| ADFTD | AD | 36 | $66.4 \pm 7.9$ | 19 | 500 | 13.5 |
| | FTD | 23 | $63.6 \pm 8.2$ | | | 12.0 |
| | HC | 29 | $67.9 \pm 5.4$ | | | 13.8 |
| CAUEEG | Dementia | 311 | $70.77 \pm 9.90$ | 19 | 200 | 13.34 |
|  | MCI | 417 |  |  |  |  |
|  | Normal | 459 |  |  |  |  |

All datasets were recorded using the 10-20 electrode setup. The common 19 electrode montage consisted of Fp1, Fp2, F7, F3, Fz, F4, F8, T3, C3, Cz, C4, T4, T5, P3, Pz, P4, T6, O1, and O2 electrodes. For the GENEEG, the electrodes T5 and T6 were removed due to technical reasons. The CAUEEG additionally had two channels associated to photic stimulation, which were removed before the preprocessing, and only 19 channels are left for further analysis.

### 2.2 Preprocessing

All EEG datasets underwent standardized pre-processing procedures to ensure comparability across sources. The pre-processing was performed using Python version 3.10.11 and the MNE-Python library (version 1.8.0) (Gramfort et al., 2013). The recordings of MCIHC and ADFTD were resampled from 256 Hz and 500 Hz, respectively, to a common sampling rate of 200 Hz, while GENEEG and CAUEEG had already been provided at 200 Hz.

A finite impulse response (FIR) band-pass filter was applied to retain frequency components between 0.5 and 45 Hz, minimizing low-frequency drifts and high-frequency noise. Then, the EEG signals were re-referenced to the common average using MNE, ensuring that each channel reflects potential differences relative to the mean scalp activity. Then, channel-wise z-score normalization was applied to standardize signal amplitudes. Finally, the continuous EEG recordings were segmented into 4-second epochs with 50% overlap, providing temporal granularity and augmenting the training dataset to help mitigate overfitting. In the CAUEEG dataset, eyes-opened and photic stimulation events were removed before the segmentation.

### 2.3 Models

This study evaluated ten DL models: MSVTNet (Liu et al., 2024), EEG-Deformer (Ding et al., 2025), EEG-Conformer (Song et al., 2023), MBSzEEGNet (Chou et al., 2024; Chang et al., 2024), Oh-CNN (Oh et al., 2019), SzHNN (Sharma and Joshi, 2022), SCCNet (Wei et al., 2019), EEGNet (Lawherm et al., 2018), DeepConvNet (Schirrmeister et al., 2017), ShallowConvNet (Schirrmeister et al., 2017). These models were selected since they demonstrated effectiveness across various resting-state EEG applications.

#### MSVTNet

MSVTNet (Liu et al., 2024) is multi-branch Vision Transformer-based model. It first extracts local spatio-temporal features at different filtered scales using CNNs. These representations are then concatenated along the feature dimension, producing multi-scale spatio-temporal tokens. The Vision Transformers capture cross-scale interaction information and global temporal correlations, resulting in more discriminative feature embeddings for classification. Additionally, auxiliary branch loss provides intermediate supervision, helping to guide and stabilize the joint learning of CNN and Transformers.

#### EEG-Deformer

EEG-Deformer (Ding et al., 2025) is a recent CNN-Transformer-based architecture. It performs a shallow CNN encoder that extracts basic temporal and spatial features from the EEG signals. These features are then processed by a Hierarchical Coarse-to-Fine Transformer, which combines parallel Transformer-based and CNN-based branches to learn both the coarse and fine-grained temporal dynamics. This model is designed to focus on neurophysiologically relevant brain regions associated with the specific cognitive task.

#### EEG-Conformer

EEG-Conformer (Song et al., 2023) combines convolution and self-attention to learn both local and global patterns in EEG data. The model begins with temporal and spatial convolutions that extract the local representation. Then, self-attention captures global temporal dependencies. Finally, a fully connected layer generates the classification output.

#### MBSzEEGNet

MBSzEEGNet (Chou et al., 2024; Chang et al., 2024) is a multi-branch model that combines Evoked Response and Rhythmic Activity extractors. Evoked Response branches focus on oscillatory activity across various frequency bands, and Rhythmic Activity branches capture spatial-spectral patterns from different brain regions. Each branch operates at its own temporal scale using distinct convolutional kernels, allowing the model to capture both high- and low-frequency features. The output of all branches is then concatenated into a single feature vector and passed through a final classifier to produce the predictions.

#### Oh-CNN

The CNN model introduced by Oh et al. (Oh et al., 2019) consists of 11 layers that combine convolutional, average pooling, and global average-pooling operations. A dropout with a rate of 0.5 was applied at the fourth and sixth layers to improve generalization. The architecture is designed to smooth feature representations and broaden the prediction range, thereby enhancing classification accuracy and enabling the model to generalize well to unseen subjects.

#### SzHNN

SzHNN, introduced in (Sharma and Joshi, 2022), is a hybrid DL model that combines CNN layers for local spatial feature extraction with LSTM layers that leverage long-term temporal dependencies for classification. By capturing both dynamic spatial and temporal characteristics, SzHNN overcomes the limitations of the purely CNN-based approaches, which overlook temporal dynamics in EEG data.

#### SCCNet

SCCNet (Wei et al., 2019) is an EEG model designed for motor imagery tasks. It introduces spatial-component-wise convolution in its initial layer to enhance feature extraction and suppress noise, followed by spatiotemporal filtering and temporal smoothing. This design prioritizes the spatial feature learning while reducing signal artifacts. Its lightweight architecture further supports better generalization and lowers the risk of overfitting.

#### EEGNet

EEGNet (Lawherm et al., 2018) is a compact CNN architecture tailored for EEG signal analysis. Its lightweight architecture performs well even with limited training data. Temporal convolution is initially applied to extract frequency components, followed by depthwise convolution to learn spatial features associated with these frequencies. Separable convolution is finally used to extract temporal features. EEGNet has been successfully applied to various EEG-related tasks and consistently demonstrates strong classification performance.

#### DeepConvNet

DeepConvNet is a deep CNN proposed by (Schirrmeister et al., 2017) for EEG decoding. It consists of multiple stacked convolutional and pooling layers designed to learn increasingly abstract representations from raw EEG signals. The first two layers perform temporal and spatial convolutions, enabling the model to extract frequency-specific activity and channel-level interactions. Subsequent convolutional blocks capture higher-level, nonlinear EEG patterns, while max-pooling layers reduce dimensionality and enhance robustness. DeepConvNet performs well on complex EEG tasks.

#### ShallowConvNet

ShallowConvNet is another CNN model introduced by (Schirrmeister et al., 2017). It is tailored for EEG decoding for motor-imagery tasks. It performs temporal convolution followed by spatial filtering, extracting band-power-related features. This structure provides strong performance with limited data and maintains good interpretability.

### 2.4 Training Strategy

The training pipeline consisted of two stages: pilot training and full training. During pilot training, a separate grid search was performed for each model–task combination to select the hyperparameters, including batch size (16, 32, 64), learning rate (1 *×* 10^−3^, 5 *×* 10^−3^, 1 *×* 10^−4^, 5 *×* 10^−4^, 1 *×* 10^−5^), and L2 regularization (0, 1 *×* 10^−2^, 1 *×* 10^−3^, 1 *×* 10^−4^). The data were divided by subject into training, validation, and test sets to prevent subject-level overlap between partitions. Each hyperparameter configuration was trained for 30 epochs on the training set, and the configuration that achieved the highest segment-level accuracy on the validation set was selected. The test set was not used for hyperparameter selection. The selected hyperparameters were subsequently used during the full training stage. The pilot split was used only for hyperparameter selection and was independent of the subsequent five-fold evaluation.

Due to overfitting, early stopping was applied. Training was terminated if the validation segment-wise accuracy did not improve for 30 consecutive epochs, and the checkpoint corresponding to the best validation performance was retained. Evaluation metrics were computed on the test set using the best validated model. To ensure the robustness of the results, the entire training and testing process was repeated 10 times using different random seeds. The seeds were generated independently for each task and stored in the GitHub repository together with the code to support reproducibility. Final performance was reported as the mean across the 10 repetitions. All models were trained on an NVIDIA RTX 4090 GPU with 24 GB of VRAM. Computational efficiency was additionally evaluated in terms of the number of trainable parameters, training time, inference time, and average peak GPU memory usage. Inference time is the GPU-synchronized wall-clock time for one forward pass over the held-out test fold, and average peak GPU memory is the peak *torch*.*cuda*.*max_memory_allocated()* reading from model instantiation through training, validation, and inference on that fold, each averaged across the 5 cross-validation folds of one representative run. The computational efficiency was reported in Appendix A, Tables 5, 7, 9, 11, 13, 15, 17, 19.

### 2.5 Model Evaluation

In this study, leave-one-fold-out cross-validation with metrics averaged across the folds was selected as the evaluation strategy. The data were separated using the StratifiedGroupKFold method implemented in the scikit-learn Python library. The identity of the subject was used as a grouping variable, ensuring that all EEG segments were assigned to the same fold and preventing subject-level data leakage between the training, validation, and test sets. Meanwhile, stratification was used to preserve the class distribution across the folds, since the datasets contained unequal numbers of the subjects in the diagnostic classes. In each iteration, three folds were used for training, one for validation, and one for testing. This strategy assessed model generalization to previously unseen subjects while reducing the imbalance across the all cross-validation folds (Kunjan et al., 2021; Lee et al., 2023).

To evaluate model performance, accuracy was computed at both the segment and subject levels. The definition of the metric depended on the evaluation granularity. Segment-wise accuracy was calculated from predictions of individual segments, where TP, TN, FP, and FN denote true positives, true negatives, false positives, and false negatives, respectively. For subject-wise evaluation, the predicted label of each subject was determined by majority voting over its segment-level predictions, with TP, TN, FP, and FN defined at the subject level. In addition to accuracy, four subject-level metrics—sensitivity, specificity, precision, and F1-score—were used to assess subject-level classification performance further (Sibilano et al., 2023).

Segment-wise accuracy provides a fine-grained assessment of model behavior across individual temporal segments. In contrast, subject-wise accuracy improves robustness by aggregating segment-level predictions, thereby mitigating the influence of noisy segments on final decisions. Sensitivity measures the model’s ability to correctly identify patients in the positive class, whereas specificity quantifies the model’s ability to correctly identify patients in the negative class. In this study, for each binary task, the first class named in the task was treated as the positive class. Precision reflects the reliability of positive predictions, and the F1-score balances precision and sensitivity, making it suitable for imbalanced datasets. Together, these metrics enable a comprehensive evaluation of model performance (Chang et al., 2024).

### 2.6 Code Availability

The source code used in this study is publicly available at https://github.com/Void97/mci-eeg. The repository also provides the random seeds and selected hyperparameters used in the experiments to facilitate reproducibility of the benchmark.

## 3 Results

This section demonstrates the results of the benchmark across the following dementia-related classification tasks: (1) MCI vs HC from GENEEG dataset; (2) MCI vs HC from MCIHC dataset; (3) AD vs HC, FTD vs HC, FTD vs AD from ADFTD dataset; (4) Dementia vs Normal, MCI vs Normal, MCI vs Dementia from CAUEEG dataset. The experimental results are presented from four complementary perspectives. First, the performance profiles of the ten deep learning models are examined across the six reported evaluation metrics for each classification task. Second, F1 scores are compared across the eight dementia-related classification tasks to examine task-specific differences in model performance. Third, the mean performance of each model across all tasks is analyzed to provide an overall comparison of the evaluated architectures. Finally, the computational requirements of the evaluated architectures are compared in terms of trainable parameters, training and inference time, and GPU memory consumption.

### 3.1 Overall Benchmark Performance

Fig. 1 visualizes the performance of the ten evaluated models. The corresponding numerical results are provided in Appendix A, Tables 4, 6, 8, 10, 12, 14, 16, and 18. The radar plots show substantial variation in performance profiles across both models and classification tasks. No architecture consistently exhibited the strongest performance across all six metrics or all diagnostic problems. In several tasks, models with comparable overall performance displayed different trade-offs among sensitivity, specificity, precision, and F1 score, demonstrating that the relative ranking of the models depended on the selected evaluation criterion. The performance profiles also differed markedly across classification tasks. Stronger overall profiles were generally observed for CAUEEG Dementia vs. Normal, ADFTD AD vs. HC, and GENEEG MCI vs. HC classification. In contrast, lower metric values were observed for several more challenging problems, particularly MCIHC MCI vs. HC, ADFTD FTD vs. HC, and CAUEEG MCI vs. Dementia classification. Two additional patterns were evident across the radar plots. First, subject-level accuracy was generally comparable to or higher than segment-level accuracy, suggesting that aggregation of epoch-level predictions frequently improved the final subject-level decision. Second, specificity was commonly higher than sensitivity, indicating a tendency for the models to identify control subjects more successfully than subjects with cognitive impairment.

**Figure 1:**
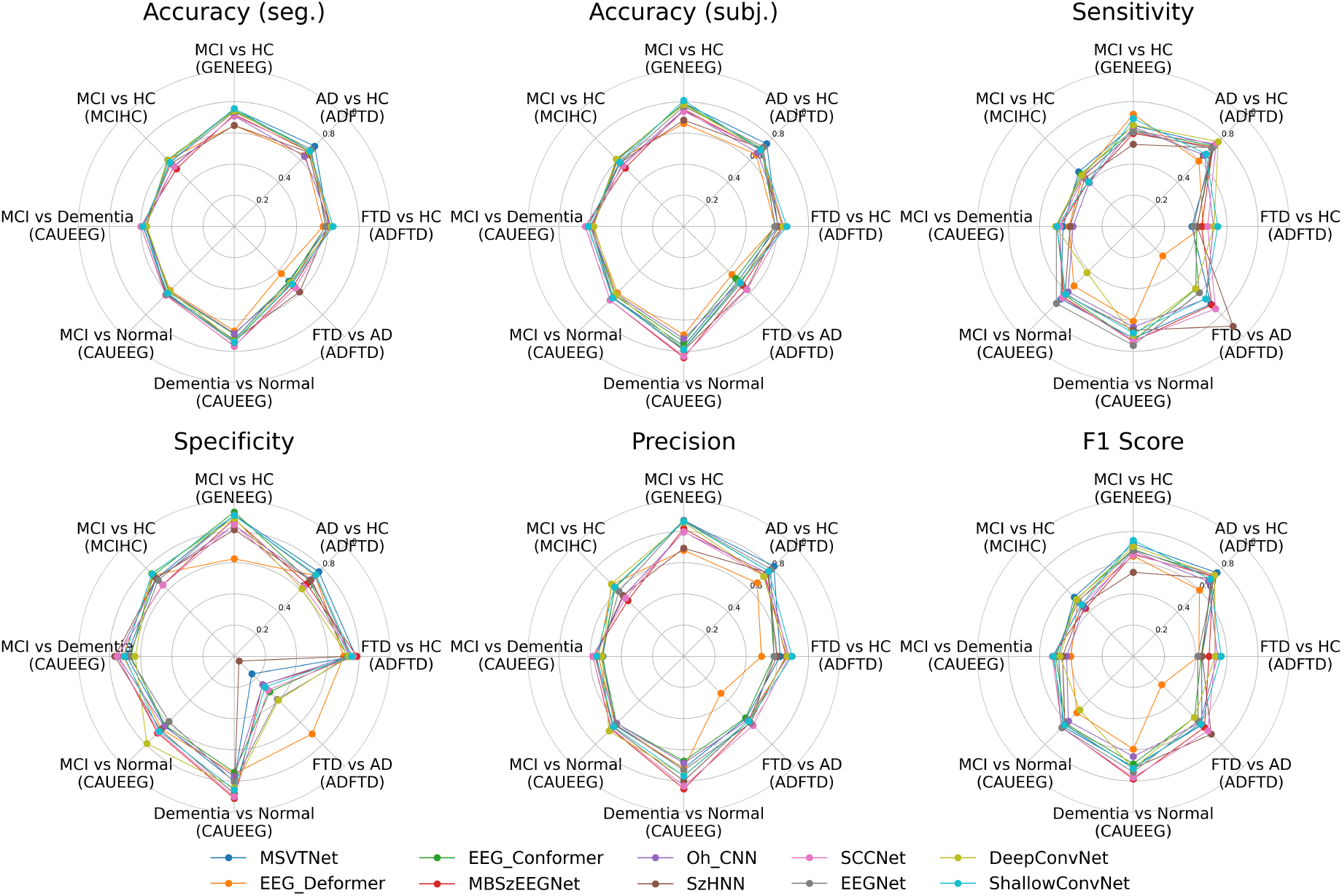
The visualization of the ten EEG models’ performance using the radar plot.

### 3.2 Mean Performance Across Classification Tasks

Table 2 summarizes the mean performance of each model across all eight classification tasks. For each architecture and evaluation metric, the task-level mean values were averaged equally across the eight classification tasks. The standard deviation reflects variability across the eight classification tasks rather than variability across repeated runs within an individual task. The aggregate results show that the leading architectures were closely matched. SCCNet obtained the highest mean subject-level accuracy (66.65%), sensitivity (61.03%), and F1 score (62.48%). In contrast, ShallowConvNet achieved the highest mean segment-level accuracy (63.90%), specificity (70.80%), and precision (68.88%). Therefore, neither model dominated across all evaluation criteria. MBSzEEGNet and MSVTNet also showed consistently competitive mean performance across several metrics. MBSzEEGNet achieved a mean subject-level accuracy of 66.04% and an F1 score of 61.20%, whereas MSVTNet achieved 65.49% and 61.28%, respectively. EEG-Deformer showed the lowest mean segment-level accuracy (58.89%), subject-level accuracy (60.24%), sensitivity (50.57%), and F1 score (49.13%).

**Table 2:**
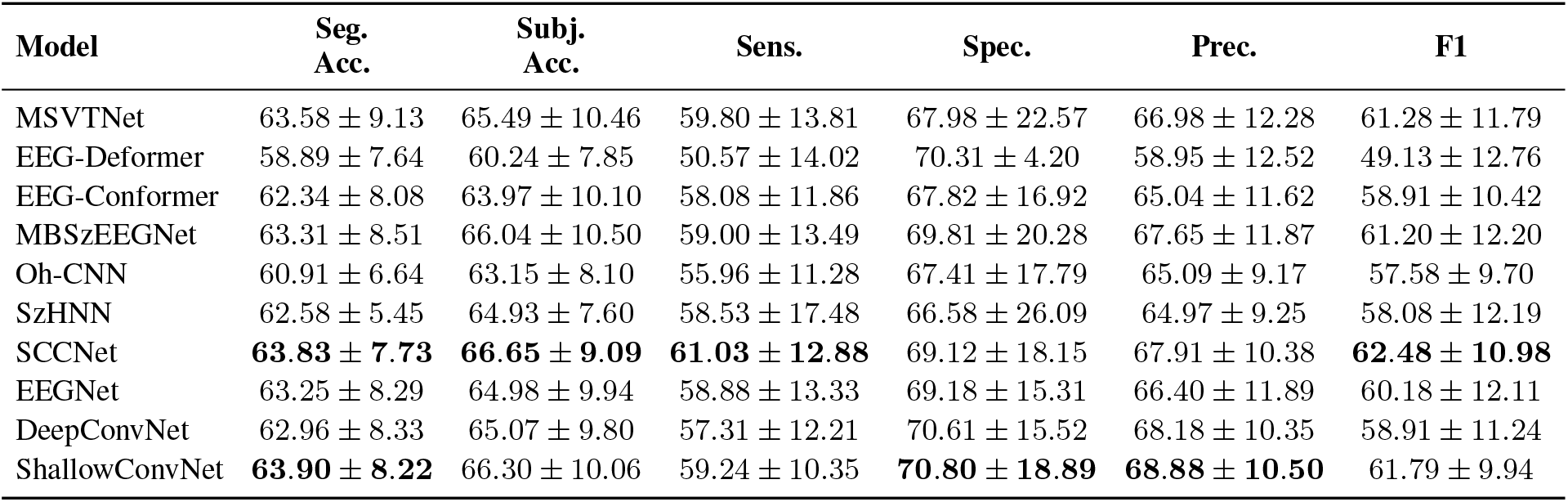
Average performance of deep learning models across all binary classification tasks. Values are reported as mean ± standard deviation across the eight classification tasks, in percentage points.

### 3.3 F1-Score Comparison Across Classification Tasks

To provide a compact comparison of model performance across the eight classification tasks, Table 3 reports the subject-level F1 score. The results show that no single architecture consistently achieved the highest F1 score. ShallowConvNet obtained the highest F1 score for GENEEG MCI vs. HC (74.42%) and ADFTD FTD vs. HC (56.35%), whereas MSVTNet performed best for MCIHC MCI vs. HC (53.58%) and ADFTD AD vs. HC (75.82%). MBSzEEGNet achieved the highest F1 scores for CAUEEG Dementia vs. Normal (78.83%) and MCI vs. Dementia (51.92%). SCCNet obtained the highest result for CAUEEG MCI vs. Normal (64.88%), while SzHNN achieved the highest F1 score for ADFTD FTD vs. AD (70.65%). Thus, five different architectures achieved the highest F1 score on at least one task, and no model ranked first on more than two of the eight classification problems. The best-performing architecture therefore depended on the specific diagnostic task.

**Table 3:**
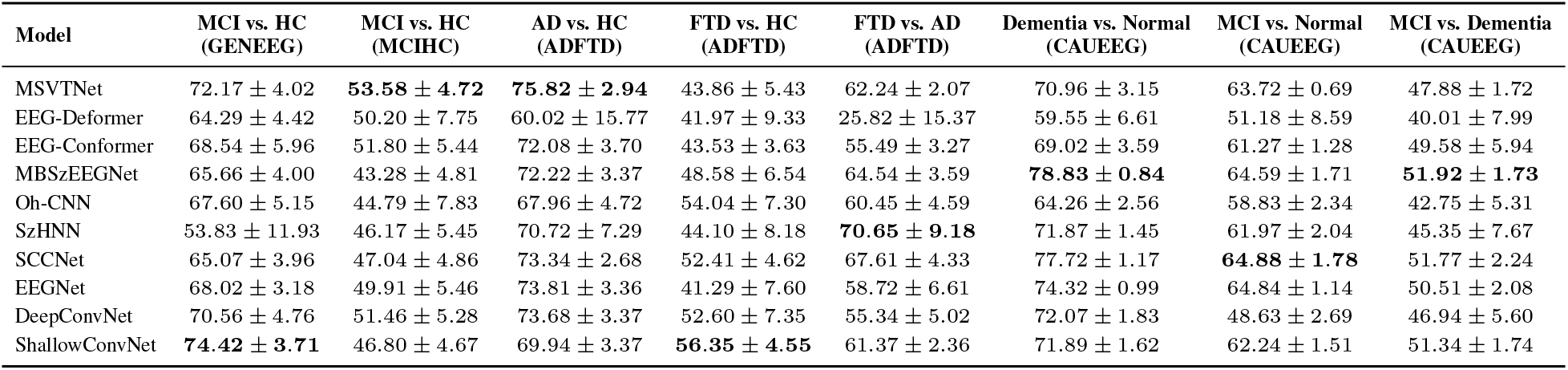
F1-score comparison of deep learning models across eight dementia-related EEG classification tasks. Values are reported as mean ± standard deviation. The best result for each task is shown in bold.

| Model | MCI vs. HC<br>(GENEEG) | MCI vs. HC<br>(MCIHC) | AD vs. HC<br>(ADFTD) | FTD vs. HC<br>(ADFTD) | FTD vs. AD<br>(ADFTD) | Dementia vs. Normal<br>(CAUEEG) | MCI vs. Normal<br>(CAUEEG) | MCI vs. Dementia<br>(CAUEEG) |
| --- | --- | --- | --- | --- | --- | --- | --- | --- |
| MSVTNet | 72.17 ± 4.02 | **53.58 ± 4.72** | **75.82 ± 2.94** | 43.86 ± 5.43 | 62.24 ± 2.07 | 70.96 ± 3.15 | 63.72 ± 0.69 | 47.88 ± 1.72 |
| EEG-Deformer | 64.29 ± 4.42 | 50.20 ± 7.75 | 60.02 ± 15.77 | 41.97 ± 9.33 | 25.82 ± 15.37 | 59.55 ± 6.61 | 51.18 ± 8.59 | 40.01 ± 7.99 |
| EEG-Conformer | 68.54 ± 5.96 | 51.80 ± 5.44 | 72.08 ± 3.70 | 43.53 ± 3.63 | 55.49 ± 3.27 | 69.02 ± 3.59 | 61.27 ± 1.28 | 49.58 ± 5.94 |
| MBSzEEGNet | 65.66 ± 4.00 | 43.28 ± 4.81 | 72.22 ± 3.37 | 48.58 ± 6.54 | 64.54 ± 3.59 | **78.83 ± 0.84** | 64.59 ± 1.71 | **51.92 ± 1.73** |
| Oh-CNN | 67.60 ± 5.15 | 44.79 ± 7.83 | 67.96 ± 4.72 | 54.04 ± 7.30 | 60.45 ± 4.59 | 64.26 ± 2.56 | 58.83 ± 2.34 | 42.75 ± 5.31 |
| SzHNN | 53.83 ± 11.93 | 46.17 ± 5.45 | 70.72 ± 7.29 | 44.10 ± 8.18 | **70.65 ± 9.18** | 71.87 ± 1.45 | 61.97 ± 2.04 | 45.35 ± 7.67 |
| SCCNet | 65.07 ± 3.96 | 47.04 ± 4.86 | 73.34 ± 2.68 | 52.41 ± 4.62 | 67.61 ± 4.33 | 77.72 ± 1.17 | **64.88 ± 1.78** | 51.77 ± 2.24 |
| EEGNet | 68.02 ± 3.18 | 49.91 ± 5.46 | 73.81 ± 3.36 | 41.29 ± 7.60 | 58.72 ± 6.61 | 74.32 ± 0.99 | 64.84 ± 1.14 | 50.51 ± 2.08 |
| DeepConvNet | 70.56 ± 4.76 | 51.46 ± 5.28 | 73.68 ± 3.37 | 52.60 ± 7.35 | 55.34 ± 5.02 | 72.07 ± 1.83 | 48.63 ± 2.69 | 46.94 ± 5.60 |
| ShallowConvNet | **74.42 ± 3.71** | 46.80 ± 4.67 | 69.94 ± 3.37 | **56.35 ± 4.55** | 61.37 ± 2.36 | 71.89 ± 1.62 | 62.24 ± 1.51 | 51.34 ± 1.74 |

Considerable differences were also observed between classification problems. The highest task-specific F1 scores were obtained for CAUEEG Dementia vs. Normal, ADFTD AD vs. HC, and GENEEG MCI vs. HC. In contrast, substantially lower values of best-performing F1-scores were observed for MCIHC MCI vs. HC, ADFTD FTD vs. HC, and CAUEEG MCI vs. Dementia. The two MCI vs. HC datasets showed particularly different performance patterns. On GENEEG, eight of the ten models achieved F1 scores above 65% (except SzHNN, whose F1-score was 53.83%), whereas all models remained below 54% on MCIHC. The highest-performing architecture also differed between the two datasets, with ShallowConvNet obtaining the best result on GENEEG and MSVTNet performing best on MCIHC.

The F1-score comparison should nevertheless be interpreted alongside the complete performance profiles shown in Fig. 1. Because F1 summarizes precision and recall but does not account for true-negative classification, a high F1 score may coexist with poor specificity. This was particularly evident for ADFTD FTD vs. AD, where SzHNN achieved the highest F1 (70.65%) score despite a strongly asymmetric sensitivity–specificity profile (90.43%, 4.30%). Accordingly, F1 was used as a compact task-wise comparison measure rather than as a standalone indicator of overall classification performance, while the complete performance profiles across all six evaluation metrics are presented in Fig. 1 and Appendix A.

### 3.4 Computational Efficiency

The evaluated architectures differed substantially in computational cost. Oh-CNN, EEGNet, and SzHNN were the smallest models, with under 10000 trainable parameters and 44-212 MB of the peak GPU memory across the tasks. In contrast, EEG-Conformer, DeepConvNet, and EEG-Deformer were the largest ones, with approximately 0.51 million, 0.58 million, and 0.69 million parameters, with EEG-Deformer consistently demonstrating the largest GPU memory consumption, which was around 13.9 GB. MSVTNet and MBSzEEGNet were in the intermediate range, with approximately 72000-95000 parameters and requiring far less memory than EEG-Conformer and EEG-Deformer.

Notably, the number of parameters was not directly proportional to the memory or runtime cost. For instance, ShallowConvNet contained only 36000-40000 parameters and required around 8.5 GB of the GPU memory, whereas MSVTNet used only around 0.71 GB despite having twice as many parameters. The runtime cost followed the same pattern. EEG-Deformer was the slowest model for training and inference, while the lightweight CNNs, such as SCCNet and Oh-CNN, remained the fastest. For instance, EEG-Deformer took 49.80 sec per training epoch on CAUEEG MCI vs Normal task, compared with 2.71 sec for Oh-CNN and 2.66 s for SCCNet. DeepConvNet showed intermediate training and inference times despite having the one of the largest parameter counts, while SzHNN, although being compact, did not always have the shortest inference time. MBSzEEGNet and MSVTNet also showed intermediate runtimes. These observations indicate that the parameter count alone is insufficient to characterize the computational efficiency of an EEG deep learning model.

Higher computational cost was not consistently associated with better classification performance. SCCNet achieved the highest subject-level accuracy and F1-score among all ten models with only 28000-31000 parameters. EEGNet also remained competitive despite its very low resource demands. MSVTNet and MBSzEEGNet combined competitive performance across several tasks with relatively lower memory consumption than the larger Transformer-based architectures. ShallowConvNet achieved strong overall performance despite its relatively high GPU memory requirement. In contrast, the larger DeepConvNet and EEG-Conformer did not demonstrate a consistent performance advantage over the compact architectures. Moreover, EEG-Deformer, despite having the largest computational cost, appeared to have the lowest subject-level accuracy and F1-score. Oh-CNN provided very low computational cost but generally lower overall performance, whereas SzHNN showed more task-dependent performance. Overall, these findings suggest that the architectural complexity and computational cost do not necessarily translate into improved dementia-related EEG classification performance. This further highlights the importance of considering both classification performance and computational efficiency for model selection.

## 4 Discussion

This study systematically compared ten deep learning architectures across eight dementia-related EEG classification tasks under a unified subject-independent evaluation protocol. Several main findings emerged. First, no single architecture consistently dominated across all tasks and evaluation metrics. Second, performance varied substantially across diagnostic problems, including between datasets representing the same nominal classification task. Third, conclusions about relative model performance depended on the selected evaluation criterion. Fourth, aggregation of epoch-level predictions generally improved subject-level classification accuracy. Fifth, the stability of the reported performance differed substantially across models and tasks. Finally, higher computational cost did not consistently correspond to better classification performance.

### 4.1 Architecture Performance

The benchmark did not identify a universally superior architecture. Five different models achieved the highest F1 score on at least one task, and no architecture ranked first on more than two of the eight classification problems. The mean results showed a similar pattern: SCCNet achieved the highest subject-level accuracy, sensitivity, and F1 score, whereas ShallowConvNet obtained the highest segment-level accuracy, specificity, and precision. MBSzEEGNet and MSVTNet also remained competitive across several tasks.

Notably, newer and more complex architectures did not consistently outperform established convolutional models. Although MSVTNet and MBSzEEGNet performed strongly on individual tasks, SCCNet and ShallowConvNet remained among the strongest models in the mean comparison. Similar findings have been reported in the standardized benchmarking of the EEG foundational models, where the greater architectural complexity did not translate uniformly into superior performance (Kastrati et al., 2025).

Performance also varied considerably between datasets and diagnostic problems. The contrast between GENEEG and MCIHC was particularly marked. Although both represent MCI vs. HC classification, the overall performance level and highest-performing architecture differed substantially. This suggests that model performance is influenced not only by architecture, but also by cohort characteristics, diagnostic procedures, acquisition conditions, and the underlying data distribution.

This interpretation is consistent with (Mishra et al., 2025), who reported dataset shifts, overlap between MCI and control EEG characteristics, and high variability in models trained on small clinical cohorts. A similar pattern was observed within CAUEEG, where Dementia vs. Normal classification was considerably stronger than MCI vs. Dementia despite the use of the same dataset and evaluation protocol.

These findings indicate that performance obtained on one dataset or task should not be treated as an intrinsic property of a model.

### 4.2 Computational Efficiency

The computational efficiency results support the findings in Section 4.1 that a more complex architecture does not necessarily lead to better classification performance. SCCNet achieved the highest mean subject-level accuracy and F1-score while remaining relatively small. Compact EEGNet also demonstrated competitive performance. Oh-CNN had the lowest computational cost among the evaluated models, although its classification performance was generally lower. SzHNN was also a lightweight model, but its inference time was not always lower than that of the larger convolutional models. In contrast, EEG-Deformer required the highest GPU memory and generally the longest training and inference time, while showing the lowest mean performance.

These differences may be partly related to the way the architectures process EEG signals. SCCNet applies spatial component-wise convolution followed by spatiotemporal filtering, while EEGNet uses depthwise and separable convolutions to extract spatial and temporal EEG features. ShallowConvNet also applies a relatively simple setup consisting of temporal and spatial convolutions, and remained one of the strongest models in the overall comparison. The architecture of DeepConvNet has deeper convolutional blocks, but it did not provide a consistent improvement in performance. Similarly, EEG-Conformer and EEG-Deformer combine CNN-based feature extraction with Transformer blocks, but their greater architectural complexity did not result in consistently better performance despite the higher computational cost. In contrast, MSVTNet and MBSzEEGNet apply multi-scale and multi-branch processing, respectively, and showed competitive results on several tasks with lower resource requirements.

It is also shown that the number of parameters alone cannot fully represent the computational cost of a model. For instance, ShallowConvNet had far fewer parameters than the Transformer-based models but still showed high GPU memory usage. DeepConvNet also had one of the largest parameter counts, but did not consistently show the longest runtime. Therefore, the computational requirements depend not only on the model size, but also on the operations performed inside the architecture during the EEG processing. These findings suggest that both classification performance and computational cost should be considered when selecting a model for EEG-based MCI and dementia classification.

### 4.3 Evaluation Metric and Model Interpretation

The six-metric profiles showed that a model performing strongly according to one criterion did not necessarily exhibit an equally strong or balanced profile across the remaining metrics.

This is particularly important for F1 score. F1 summarizes precision and sensitivity through their harmonic mean (Sibilano et al., 2023; Lipton et al., 2014), but it does not incorporate true-negative classification and therefore cannot describe specificity or the complete error profile (Powers, 2011). The ADFTD FTD vs. AD task provides a clear example: SzHNN achieved the highest F1 score (70.65%), but this result was accompanied by highly asymmetric sensitivity and specificity of 90.43% and 4.30%, respectively.

The results therefore support the complementary presentation adopted in this study. Table 3 provides a compact comparison of F1 scores across tasks, whereas Fig. 1 and Appendix A provide the broader performance profiles needed to interpret them.

A consistent difference between sensitivity and specificity was also observed in the mean results, with specificity exceeding sensitivity for every architecture. For patient-versus-control tasks, this indicates a greater tendency to correctly identify controls than affected individuals. These findings reinforce the importance of interpreting complementary metrics rather than relying on a single summary measure.

### 4.4 Subject-Level Evaluation

All ten architectures achieved higher mean subject-level accuracy than segment-level accuracy. This suggests that aggregating predictions across multiple EEG epochs generally improved the final classification of unseen subjects.

This distinction is important in clinical EEG applications, where the final prediction concerns an individual rather than a single short segment. Majority voting can reduce the influence of transient or unrepresentative epochs by combining predictions across the recording. The importance of subject-based evaluation has also been emphasized in large-scale EEG studies showing that data-partitioning strategy can substantially affect reported performance (Del Pup et al., 2025). Subject-level majority voting has likewise been used in recent EEG-based dementia research (Chang et al., 2024; Wang et al., 2025).

The Appendix A results additionally showed that performance variability differed across models and tasks. Some model–task combinations exhibited large standard deviations across the ten repetitions, whereas others were considerably more stable. This highlights the importance of repeated evaluation, particularly for smaller or more difficult clinical EEG tasks.

Although majority voting improved mean performance, it remains a simple aggregation strategy that assigns equal importance to all segments. Future work could investigate confidence-weighted or learnable subject-level aggregation.

Overall, the benchmark demonstrates that conclusions about model superiority depend on the task, dataset, evaluation metric, and aggregation level. Under a common experimental protocol, no single architecture consistently dominated.

### 4.5 Limitations

Several limitations should be considered when interpreting the results. First, although four datasets and eight classification tasks were included, each dataset was evaluated independently. The study therefore assesses within-dataset subject-independent performance and does not examine cross-dataset transfer or generalization to recordings acquired at previously unseen clinical sites. Second, the included datasets differ in cohort composition, diagnostic procedures, recording duration, channel configuration, and acquisition system. Although the unified protocol improves methodological comparability, it cannot remove these underlying differences or fully control for potential demographic and clinical confounders.

Third, applying the same preprocessing framework and hyperparameter search space to all models introduces a trade-off between fair comparison and model-specific optimization. The benchmark therefore measures performance under a common protocol rather than the maximum performance achievable by each architecture. In addition, hyperparameters and checkpoints were selected using segment-level validation accuracy, whereas the final prediction was evaluated at the subject level. The reported standard deviations describe variability across repeated experiments, but formal statistical comparisons between models were not performed. Therefore, small differences between models with similar mean performance should be interpreted cautiously.

Fourth, although ten deep learning architectures were evaluated, they do not represent all existing EEG classification models. The computational efficiency results are also dependent on the hardware and implementation used in this study, and the absolute training time, inference time, and GPU memory usage may differ on other systems. Finally, the present study focuses on predictive performance and does not investigate the neurophysiological features underlying individual model decisions. Future work should examine whether the strongest-performing architectures rely on clinically meaningful EEG characteristics and whether these patterns remain stable across independent cohorts.

## 5 Conclusion

This study presented a reproducible benchmark of ten deep learning architectures for subject-independent EEG-based MCI and dementia classification across four datasets and eight binary classification tasks. Using a unified preprocessing, training, and evaluation protocol, the benchmark enabled direct comparison of model performance under consistent experimental conditions. SCCNet and ShallowConvNet achieved the strongest mean performance on different evaluation criteria, while the best-performing model varied across classification problems. Subject-level aggregation improved mean accuracy for all evaluated architectures. Higher computational cost did not consistently correspond to better classification performance, with several relatively compact architectures remaining competitive with substantially larger models. These findings demonstrate that EEG dementia classification performance is strongly influenced by the dataset, diagnostic task, evaluation metric, prediction granularity, and model efficiency. The proposed benchmark therefore provides a reproducible framework for more reliable comparison of EEG deep learning models and highlights the need for subject-independent evaluation that considers both complementary performance metrics and computational requirements.

## A Detailed Performance and Computational Efficiency Results

Appendix A provides the complete task-level performance and computational efficiency results for all ten evaluated models. For each classification task, performance is reported in terms of segment-level accuracy, subject-level accuracy, sensitivity, specificity, precision, and F1-score. Performance values are presented as mean ± standard deviation across the ten repeated experiments and are reported in percentage points. The corresponding computational efficiency tables report the number of trainable parameters, training time per epoch, inference time, and average peak GPU memory usage. As defined in Section 2.5, the first class named in each binary classification task was treated as the positive class.

### A.1 GENEEG dataset

Tables 4 and 5 report the classification performance and computational efficiency, respectively, for the MCI vs. HC task on the GENEEG dataset.

**Table 4:**
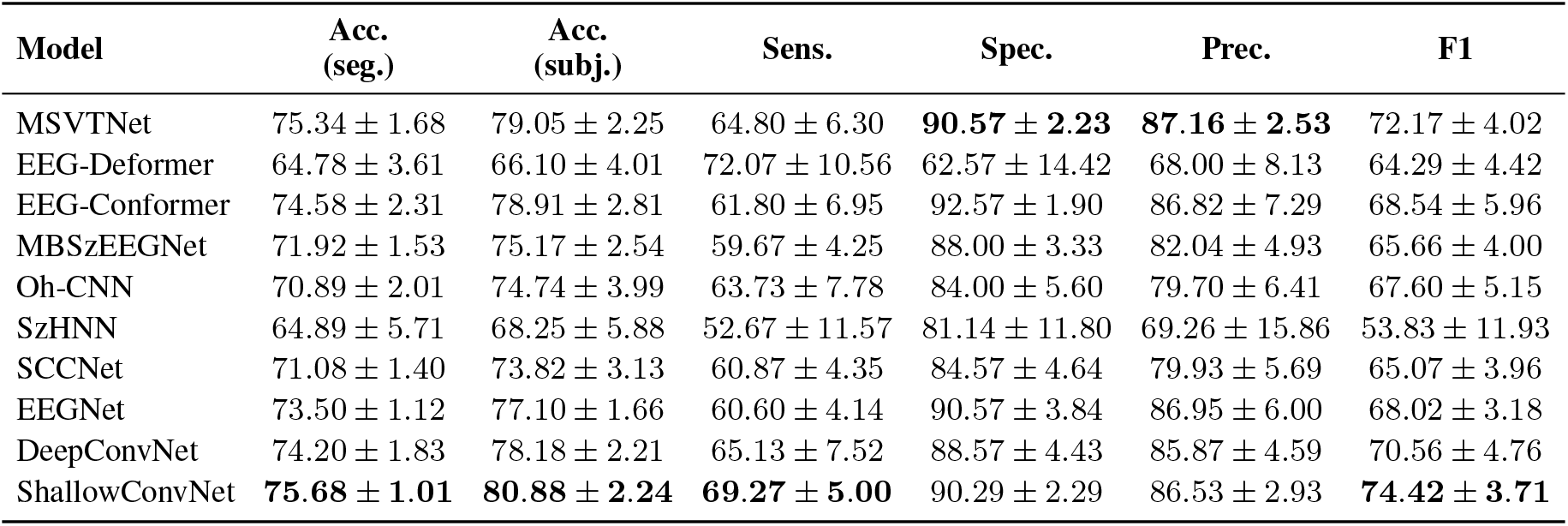
Performance comparison of deep learning models on the dataset MCI vs. HC classification task. Values are reported as mean ± standard deviation in percentage points.

| Model | Acc.<br>(seg.) | Acc.<br>(subj.) | Sens. | Spec. | Prec. | F1 |
| --- | --- | --- | --- | --- | --- | --- |
| MSVTNet | 75.34 ± 1.68 | 79.05 ± 2.25 | 64.80 ± 6.30 | **90.57 ± 2.23** | **87.16 ± 2.53** | 72.17 ± 4.02 |
| EEG-Deformer | 64.78 ± 3.61 | 66.10 ± 4.01 | 72.07 ± 10.56 | 62.57 ± 14.42 | 68.00 ± 8.13 | 64.29 ± 4.42 |
| EEG-Conformer | 74.58 ± 2.31 | 78.91 ± 2.81 | 61.80 ± 6.95 | 92.57 ± 1.90 | 86.82 ± 7.29 | 68.54 ± 5.96 |
| MBSzEEGNet | 71.92 ± 1.53 | 75.17 ± 2.54 | 59.67 ± 4.25 | 88.00 ± 3.33 | 82.04 ± 4.93 | 65.66 ± 4.00 |
| Oh-CNN | 70.89 ± 2.01 | 74.74 ± 3.99 | 63.73 ± 7.78 | 84.00 ± 5.60 | 79.70 ± 6.41 | 67.60 ± 5.15 |
| SzHNN | 64.89 ± 5.71 | 68.25 ± 5.88 | 52.67 ± 11.57 | 81.14 ± 11.80 | 69.26 ± 15.86 | 53.83 ± 11.93 |
| SCCNet | 71.08 ± 1.40 | 73.82 ± 3.13 | 60.87 ± 4.35 | 84.57 ± 4.64 | 79.93 ± 5.69 | 65.07 ± 3.96 |
| EEGNet | 73.50 ± 1.12 | 77.10 ± 1.66 | 60.60 ± 4.14 | 90.57 ± 3.84 | 86.95 ± 6.00 | 68.02 ± 3.18 |
| DeepConvNet | 74.20 ± 1.83 | 78.18 ± 2.21 | 65.13 ± 7.52 | 88.57 ± 4.43 | 85.87 ± 4.59 | 70.56 ± 4.76 |
| ShallowConvNet | **75.68 ± 1.01** | **80.88 ± 2.24** | **69.27 ± 5.00** | 90.29 ± 2.29 | 86.53 ± 2.93 | **74.42 ± 3.71** |

**Table 5:**
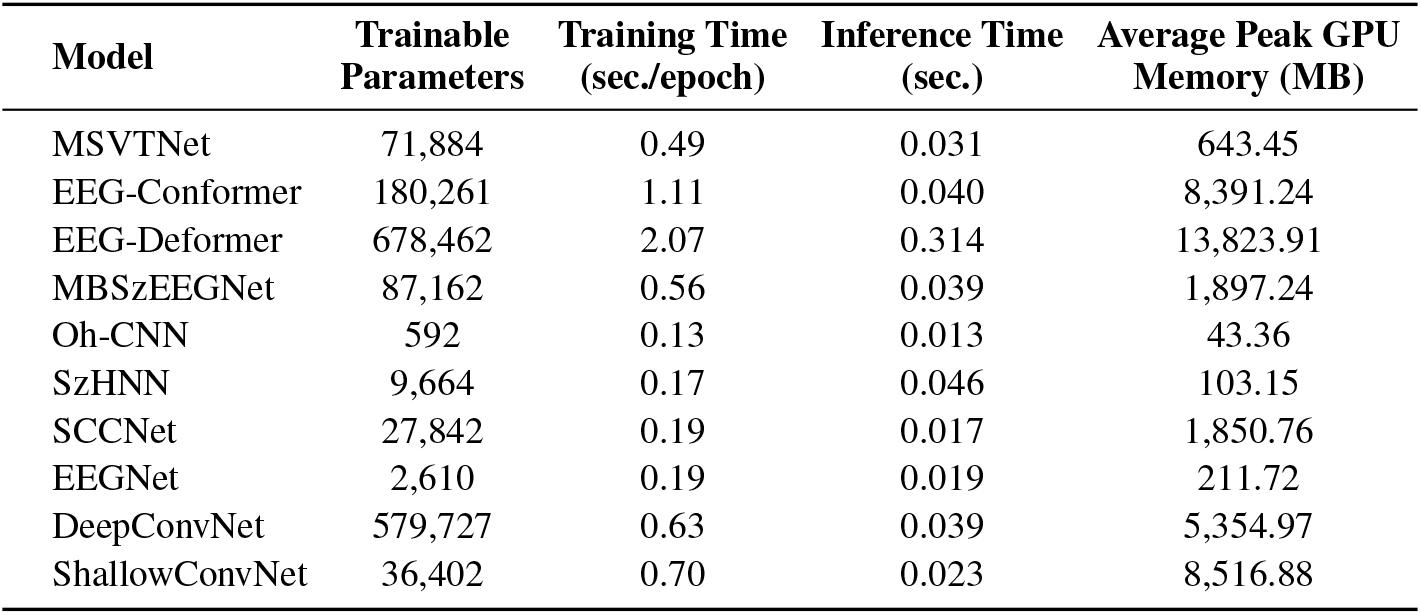
Computational efficiency of the evaluated models for the MCI vs. HC classification task.

| Model | Trainable<br>Parameters | Training Time<br>(sec/epoch) | Inference Time<br>(sec.) | Average Peak GPU<br>Memory (MB) |
| --- | --- | --- | --- | --- |
| MSVTNet | 71,884 | 0.49 | 0.031 | 643.45 |
| EEG-Conformer | 180,261 | 1.11 | 0.040 | 8,391.24 |
| EEG-Deformer | 678,462 | 2.07 | 0.314 | 13,823.91 |
| MBSzEEGNet | 87,162 | 0.56 | 0.039 | 1,897.24 |
| Oh-CNN | 592 | 0.13 | 0.013 | 43.36 |
| SzHNN | 9,664 | 0.17 | 0.046 | 103.15 |
| SCCNet | 27,842 | 0.19 | 0.017 | 1,850.76 |
| EEGNet | 2,610 | 0.19 | 0.019 | 211.72 |
| DeepConvNet | 579,727 | 0.63 | 0.039 | 5,354.97 |
| ShallowConvNet | 36,402 | 0.70 | 0.023 | 8,516.88 |

### A.2 MCIHC dataset

Tables 6 and 7 report the classification performance and computational efficiency, respectively, for the MCI vs. HC task on the MCIHC dataset.

**Table 6:** Performance comparison of deep learning models on the dataset MCI vs. HC classification task. Values are reported as mean ± standard deviation in percentage points.

| Model | Seg. Acc. | Subj. Acc. | Sens. | Spec. | Prec. | F1 |
| --- | --- | --- | --- | --- | --- | --- |
| MSVTNet | 59.77 ± 2.88 | 60.45 ± 3.47 | **49.53 ± 5.70** | 70.62 ± 4.70 | 62.41 ± 5.29 | **53.58 ± 4.72** |
| EEG-Deformer | 60.19 ± 1.74 | 59.90 ± 2.75 | 45.80 ± 12.63 | 73.52 ± 9.04 | 65.02 ± 4.06 | 50.20 ± 7.75 |
| EEG-Conformer | **60.40 ± 2.86** | 60.88 ± 3.26 | 47.60 ± 5.72 | 73.33 ± 3.87 | 62.28 ± 4.68 | 51.80 ± 5.44 |
| MBSzEEGNet | 52.31 ± 2.83 | 53.03 ± 3.78 | 40.53 ± 5.74 | 64.81 ± 5.97 | 50.60 ± 6.80 | 43.28 ± 4.81 |
| Oh-CNN | 54.79 ± 3.89 | 55.98 ± 5.37 | 39.87 ± 7.42 | 70.52 ± 7.16 | 54.62 ± 11.25 | 44.79 ± 7.83 |
| SzHNN | 57.23 ± 3.32 | 57.74 ± 4.80 | 43.93 ± 5.67 | 70.76 ± 7.31 | 55.12 ± 11.23 | 46.17 ± 5.45 |
| SCCNet | 54.35 ± 2.74 | 55.14 ± 2.63 | 45.20 ± 5.69 | 64.67 ± 4.53 | 52.83 ± 5.35 | 47.04 ± 4.86 |
| EEGNet | 58.15 ± 3.92 | 57.73 ± 4.91 | 45.80 ± 5.13 | 68.91 ± 6.80 | 58.87 ± 6.87 | 49.91 ± 5.46 |
| DeepConvNet | 60.15 ± 1.95 | **61.19 ± 4.06** | 46.60 ± 7.21 | **75.14 ± 6.72** | **65.65 ± 7.10** | 51.46 ± 5.28 |
| ShallowConvNet | 58.35 ± 1.96 | 58.35 ± 2.57 | 40.60 ± 6.15 | 74.76 ± 6.48 | 62.77 ± 7.19 | 46.80 ± 4.67 |

**Table 7:** Computational efficiency of the evaluated models for the MCI vs. HC classification task.

| Model | Trainable Parameters | Training Time (sec./epoch) | Inference Time (sec.) | Average Peak GPU Memory (MB) |
| --- | --- | --- | --- | --- |
| MSVTNet | 72,028 | 6.20 | 0.314 | 707.69 |
| EEG-Deformer | 686,654 | 12.52 | 1.881 | 13,908.05 |
| EEG-Conformer | 240,421 | 10.88 | 0.242 | 8,439.41 |
| MBSzEEGNet | 94,582 | 2.59 | 0.242 | 2,103.57 |
| Oh-CNN | 622 | 0.55 | 0.074 | 44.05 |
| SzHNN | 9,814 | 0.86 | 0.258 | 100.84 |
| SCCNet | 31,120 | 0.80 | 0.100 | 2,054.48 |
| EEGNet | 2,418 | 1.34 | 0.115 | 186.25 |
| DeepConvNet | 580,977 | 4.04 | 0.230 | 5,388.14 |
| ShallowConvNet | 39,602 | 4.48 | 0.143 | 8,578.26 |

### A.3 ADFTD

Tables 8–13 provide the detailed results for the three binary classification tasks constructed from the ADFTD dataset: AD vs. HC, FTD vs. HC, and FTD vs. AD.

**Table 8:** Performance comparison of deep learning models on the AD vs. HC classification task. Values are reported as mean ± standard deviation in percentage points.

| Model | Seg. Acc. | Subj. Acc. | Sens. | Spec. | Prec. | F1 |
| --- | --- | --- | --- | --- | --- | --- |
| MSVTNet | **72.90 ± 1.07** | **75.08 ± 3.14** | 73.89 ± 4.44 | **76.60 ± 7.78** | **81.70 ± 6.19** | **75.82 ± 2.94** |
| EEG-Deformer | 64.65 ± 6.19 | 65.54 ± 7.60 | 59.39 ± 15.00 | 73.20 ± 11.80 | 66.54 ± 19.49 | 60.02 ± 15.77 |
| EEG-Conformer | 69.82 ± 2.10 | 70.61 ± 3.19 | 73.00 ± 6.72 | 68.33 ± 9.67 | 77.04 ± 6.25 | 72.08 ± 3.70 |
| MBSzEEGNet | 67.96 ± 1.29 | 69.54 ± 3.06 | 72.11 ± 5.12 | 65.73 ± 4.57 | 73.56 ± 2.77 | 72.22 ± 3.37 |
| Oh-CNN | 63.68 ± 2.05 | 66.92 ± 4.73 | 64.61 ± 6.36 | 69.73 ± 9.49 | 75.59 ± 6.13 | 67.96 ± 4.72 |
| SzHNN | 67.31 ± 2.91 | 70.77 ± 5.59 | 71.43 ± 9.18 | 69.00 ± 15.08 | 75.98 ± 10.90 | 70.72 ± 7.29 |
| SCCNet | 68.75 ± 0.96 | 69.84 ± 2.85 | 75.00 ± 2.86 | 63.00 ± 4.96 | 72.73 ± 3.37 | 73.34 ± 2.68 |
| EEGNet | 69.83 ± 1.02 | 72.15 ± 2.52 | 70.96 ± 5.51 | 73.33 ± 3.71 | 78.97 ± 3.79 | 73.81 ± 3.36 |
| DeepConvNet | 67.98 ± 2.10 | 69.69 ± 4.30 | **76.96 ± 4.16** | 61.33 ± 9.47 | 72.36 ± 5.43 | 73.68 ± 3.37 |
| ShallowConvNet | 68.54 ± 1.91 | 69.54 ± 2.90 | 65.82 ± 4.13 | 74.07 ± 3.97 | 77.01 ± 2.94 | 69.94 ± 3.37 |

**Table 9:** Computational efficiency of the evaluated models for the AD vs. HC classification task.

| Model | Trainable Parameters | Training Time (sec./epoch) | Inference Time (sec.) | Average Peak GPU Memory (MB) |
| --- | --- | --- | --- | --- |
| MSVTNet | 72,028 | 2.83 | 0.151 | 707.51 |
| EEG-Deformer | 686,654 | 6.57 | 0.888 | 13,913.60 |
| EEG-Conformer | 511,285 | 5.02 | 0.116 | 8,475.82 |
| MBSzEEGNet | 94,582 | 1.16 | 0.116 | 2,102.75 |
| Oh-CNN | 622 | 0.64 | 0.035 | 44.07 |
| SzHNN | 9,814 | 0.77 | 0.229 | 102.79 |
| SCCNet | 31,120 | 0.43 | 0.048 | 2,054.72 |
| EEGNet | 2,418 | 0.38 | 0.055 | 185.36 |
| DeepConvNet | 580,977 | 1.85 | 0.101 | 5,393.19 |
| ShallowConvNet | 39,602 | 2.08 | 0.230 | 8,578.56 |

**Table 10:** Performance comparison of deep learning models on the FTD vs. HC classification task. Values are reported as mean ± standard deviation in percentage points.

| Model | Seg. Acc. | Subj. Acc. | Sens. | Spec. | Prec. | F1 |
| --- | --- | --- | --- | --- | --- | --- |
| MSVTNet | 60.55 ± 2.41 | 60.29 ± 2.69 | 37.50 ± 6.45 | 77.93 ± 6.93 | 61.76 ± 6.58 | 43.86 ± 5.43 |
| EEG-Deformer | 56.88 ± 4.84 | 58.67 ± 4.76 | 44.90 ± 12.95 | 70.13 ± 13.64 | 49.83 ± 11.15 | 41.97 ± 9.33 |
| EEG-Conformer | 58.64 ± 3.21 | 58.64 ± 4.03 | 40.80 ± 5.42 | 72.60 ± 8.36 | 57.78 ± 5.53 | 43.53 ± 3.63 |
| MBSzEEGNet | 61.80 ± 3.75 | 63.22 ± 5.38 | 43.80 ± 6.35 | **78.67 ± 6.94** | 65.61 ± 8.70 | 48.58 ± 6.54 |
| Oh-CNN | 61.75 ± 2.77 | 64.27 ± 4.46 | 51.50 ± 8.49 | 73.93 ± 5.07 | 66.76 ± 6.23 | 54.04 ± 7.30 |
| SzHNN | 59.85 ± 1.59 | 60.64 ± 4.11 | 41.00 ± 10.79 | 75.87 ± 7.31 | 59.68 ± 8.88 | 44.10 ± 8.18 |
| SCCNet | 62.06 ± 1.75 | 63.94 ± 3.36 | 47.70 ± 4.56 | 76.33 ± 5.22 | 67.87 ± 6.23 | 52.41 ± 4.62 |
| EEGNet | 58.55 ± 3.38 | 58.33 ± 4.47 | 38.60 ± 9.95 | 74.33 ± 10.01 | 57.86 ± 10.19 | 41.29 ± 7.60 |
| DeepConvNet | 62.15 ± 2.62 | 63.49 ± 4.02 | 51.70 ± 8.98 | 73.00 ± 5.60 | 66.13 ± 6.21 | 52.60 ± 7.35 |
| ShallowConvNet | **63.29 ± 2.10** | **66.05 ± 3.30** | **54.10 ± 5.96** | 75.20 ± 3.36 | **69.37 ± 3.96** | **56.35 ± 4.55** |

**Table 11:** Computational efficiency of the evaluated models for the FTD vs. HC classification task.

| Model | Trainable Parameters | Training Time (sec./epoch) | Inference Time (sec.) | Average Peak GPU Memory (MB) |
| --- | --- | --- | --- | --- |
| MSVTNet | 72,028 | 2.19 | 0.116 | 707.51 |
| EEG-Deformer | 686,654 | 5.07 | 0.681 | 13,913.60 |
| EEG-Conformer | 511,285 | 3.90 | 0.089 | 8,475.48 |
| MBSzEEGNet | 94,582 | 1.26 | 0.089 | 2,102.73 |
| Oh-CNN | 622 | 0.30 | 0.028 | 44.07 |
| SzHNN | 9,814 | 0.23 | 0.093 | 103.91 |
| SCCNet | 31,120 | 0.28 | 0.038 | 2,054.72 |
| EEGNet | 2,418 | 0.49 | 0.042 | 186.28 |
| DeepConvNet | 580,977 | 1.40 | 0.094 | 5,392.52 |
| ShallowConvNet | 39,602 | 1.61 | 0.053 | 8,579.35 |

**Table 12:** Performance comparison of deep learning models on the FTD vs. AD classification task. Values are reported as mean ± standard deviation in percentage points.

| Model | Seg. Acc. | Subj. Acc. | Sens. | Spec. | Prec. | F1 |
| --- | --- | --- | --- | --- | --- | --- |
| MSVTNet | 49.43 ± 2.05 | 49.22 ± 2.00 | 70.36 ± 3.42 | 16.00 ± 4.40 | 56.39 ± 1.47 | 62.24 ± 2.07 |
| EEG-Deformer | 42.61 ± 5.01 | 43.59 ± 4.91 | 26.54 ± 16.26 | **70.50 ± 14.90** | 33.55 ± 21.87 | 25.82 ± 15.37 |
| EEG-Conformer | 50.25 ± 1.27 | 47.16 ± 2.30 | 56.71 ± 4.76 | 32.20 ± 6.55 | 55.91 ± 2.52 | 55.49 ± 3.27 |
| MBSzEEGNet | 53.83 ± 2.53 | 53.30 ± 3.14 | 71.14 ± 8.37 | 25.70 ± 10.79 | 60.05 ± 2.57 | 64.54 ± 3.59 |
| Oh-CNN | 51.60 ± 2.83 | 50.42 ± 3.62 | 65.68 ± 9.03 | 25.90 ± 12.09 | 57.99 ± 3.32 | 60.45 ± 4.59 |
| SzHNN | **59.24 ± 7.15** | 56.87 ± 7.26 | **90.43 ± 16.71** | 4.30 ± 7.77 | 58.65 ± 4.47 | **70.65 ± 9.18** |
| SCCNet | 55.67 ± 2.23 | **57.25 ± 4.80** | 74.54 ± 6.33 | 30.60 ± 6.48 | **62.76 ± 3.56** | 67.61 ± 4.33 |
| EEGNet | 51.82 ± 3.28 | 51.71 ± 5.80 | 59.86 ± 9.04 | 39.10 ± 8.60 | 60.54 ± 5.24 | 58.72 ± 6.61 |
| DeepConvNet | 50.85 ± 1.82 | 50.41 ± 2.43 | 56.39 ± 6.70 | 39.90 ± 9.19 | 58.80 ± 4.89 | 55.34 ± 5.02 |
| ShallowConvNet | 52.24 ± 1.41 | 50.78 ± 2.73 | 65.61 ± 2.76 | 27.90 ± 4.74 | 58.86 ± 2.04 | 61.37 ± 2.36 |

**Table 13:** Computational efficiency of the evaluated models for the FTD vs. AD classification task.

| Model | Trainable Parameters | Training Time (sec./epoch) | Inference Time (sec.) | Average Peak GPU Memory (MB) |
| --- | --- | --- | --- | --- |
| MSVTNet | 72,028 | 1.38 | 0.1303 | 707.69 |
| EEG-Deformer | 686,654 | 5.03 | 0.7639 | 13,908.05 |
| EEG-Conformer | 511,285 | 4.33 | 0.0993 | 8,439.41 |
| MBSzEEGNet | 94,582 | 0.92 | 0.0993 | 2,103.57 |
| Oh-CNN | 622 | 0.33 | 0.0315 | 44.05 |
| SzHNN | 9,814 | 0.63 | 0.1010 | 100.84 |
| SCCNet | 31,120 | 0.33 | 0.0414 | 2,054.47 |
| EEGNet | 2,418 | 0.41 | 0.0473 | 186.25 |
| DeepConvNet | 580,977 | 1.58 | 0.0941 | 5,388.14 |
| ShallowConvNet | 39,602 | 1.81 | 0.0588 | 8,578.26 |

### A.4 CAUEEG dataset

Tables 14–19 provide the detailed results for the three binary classification tasks constructed from the CAUEEG dataset: Dementia vs. Normal, MCI vs. Normal, and MCI vs. Dementia.

**Table 14:** Performance comparison of deep learning models on the Dementia vs. Normal classification task. Values are reported as mean ± standard deviation in percentage points.

| Model | Seg. Acc. | Subj. Acc. | Sens. | Spec. | Prec. | F1 |
| --- | --- | --- | --- | --- | --- | --- |
| MSVTNet | 72.65 ± 0.76 | 76.75 ± 1.02 | 71.69 ± 7.45 | 80.16 ± 4.32 | 71.94 ± 3.94 | 70.96 ± 3.15 |
| EEG-Deformer | 66.98 ± 7.20 | 69.50 ± 9.28 | 60.78 ± 9.55 | 75.36 ± 20.34 | 71.83 ± 12.68 | 59.55 ± 6.61 |
| EEG-Conformer | 69.07 ± 1.44 | 73.98 ± 1.68 | 73.42 ± 8.30 | 74.32 ± 5.17 | 67.24 ± 2.54 | 69.02 ± 3.59 |
| MBSzEEGNet | **76.86 ± 0.30** | **84.15 ± 0.54** | 73.76 ± 1.68 | **91.15 ± 0.79** | **85.06 ± 0.96** | **78.83 ± 0.84** |
| Oh-CNN | 68.55 ± 0.80 | 72.03 ± 1.48 | 64.48 ± 4.99 | 77.13 ± 3.87 | 68.58 ± 1.92 | 64.26 ± 2.56 |
| SzHNN | 73.09 ± 1.23 | 79.43 ± 0.47 | 66.49 ± 3.74 | 88.14 ± 2.47 | 80.02 ± 1.98 | 71.87 ± 1.45 |
| SCCNet | 76.60 ± 0.37 | 83.26 ± 0.73 | 73.17 ± 1.92 | 90.06 ± 0.88 | 83.27 ± 1.19 | 77.72 ± 1.17 |
| EEGNet | 74.43 ± 0.66 | 78.87 ± 1.12 | **76.27 ± 1.67** | 80.60 ± 2.47 | 73.14 ± 1.96 | 74.32 ± 0.99 |
| DeepConvNet | 73.60 ± 0.90 | 78.43 ± 1.34 | 69.86 ± 4.48 | 84.22 ± 3.87 | 76.53 ± 3.48 | 72.07 ± 1.83 |
| ShallowConvNet | 74.12 ± 0.49 | 78.70 ± 0.92 | 68.26 ± 2.74 | 85.73 ± 1.65 | 76.78 ± 1.79 | 71.89 ± 1.62 |

**Table 15:** Computational efficiency of the evaluated models for the Dementia vs. Normal classification task.

| Model | Trainable Parameters | Training Time (sec./epoch) | Inference Time (sec.) | Average Peak GPU Memory (MB) |
| --- | --- | --- | --- | --- |
| MSVTNet | 72,028 | 13.02 | 0.926 | 707.51 |
| EEG-Deformer | 686,654 | 44.25 | 5.612 | 13,913.60 |
| EEG-Conformer | 240,421 | 26.73 | 0.707 | 8,471.12 |
| MBSzEEGNet | 94,582 | 7.26 | 0.704 | 2,102.85 |
| Oh-CNN | 622 | 1.62 | 0.215 | 44.07 |
| SzHNN | 9,814 | 4.70 | 0.894 | 103.50 |
| SCCNet | 31,120 | 3.17 | 0.294 | 2,055.47 |
| EEGNet | 2,418 | 2.89 | 0.338 | 185.36 |
| DeepConvNet | 580,977 | 11.70 | 0.675 | 5,392.54 |
| ShallowConvNet | 39,602 | 13.07 | 0.420 | 8,579.35 |

**Table 16:** Performance comparison of deep learning models on the MCI vs. Normal classification task. Values are reported as mean ± standard deviation in percentage points.

| Model | Seg. Acc. | Subj. Acc. | Sens. | Spec. | Prec. | F1 |
| --- | --- | --- | --- | --- | --- | --- |
| MSVTNet | 61.28 ± 0.55 | 64.32 ± 0.95 | 65.69 ± 1.84 | 63.05 ± 2.91 | 62.56 ± 1.42 | 63.72 ± 0.69 |
| EEG-Deformer | 58.13 ± 3.05 | 60.21 ± 3.69 | 53.84 ± 13.32 | 66.00 ± 13.40 | 62.04 ± 9.38 | 51.18 ± 8.59 |
| EEG-Conformer | 59.42 ± 0.71 | 62.59 ± 0.93 | 62.38 ± 2.97 | 62.77 ± 3.19 | 60.91 ± 1.25 | 61.27 ± 1.28 |
| MBSzEEGNet | 62.02 ± 0.36 | **66.82 ± 0.58** | 63.84 ± 3.70 | 69.55 ± 2.55 | 65.92 ± 0.59 | 64.59 ± 1.71 |
| Oh-CNN | 58.76 ± 0.65 | 61.48 ± 1.40 | 59.18 ± 4.81 | 63.56 ± 4.90 | 60.84 ± 2.21 | 58.83 ± 2.34 |
| SzHNN | 60.83 ± 0.91 | 64.47 ± 1.52 | 61.82 ± 6.19 | 66.87 ± 7.48 | 64.29 ± 2.26 | 61.97 ± 2.04 |
| SCCNet | **62.30 ± 0.58** | 66.80 ± 1.04 | 64.68 ± 3.04 | 68.73 ± 1.56 | 65.54 ± 0.87 | **64.88 ± 1.78** |
| EEGNet | 61.53 ± 0.54 | 64.18 ± 1.33 | **69.71 ± 4.14** | 59.11 ± 5.78 | 61.78 ± 2.34 | 64.84 ± 1.14 |
| DeepConvNet | 58.63 ± 0.83 | 61.42 ± 1.14 | 41.90 ± 4.44 | **79.17 ± 4.01** | **67.77 ± 2.24** | 48.63 ± 2.69 |
| ShallowConvNet | 60.48 ± 0.43 | 64.78 ± 0.61 | 61.33 ± 3.39 | 67.96 ± 3.40 | 63.83 ± 1.54 | 62.24 ± 1.51 |

**Table 17:**
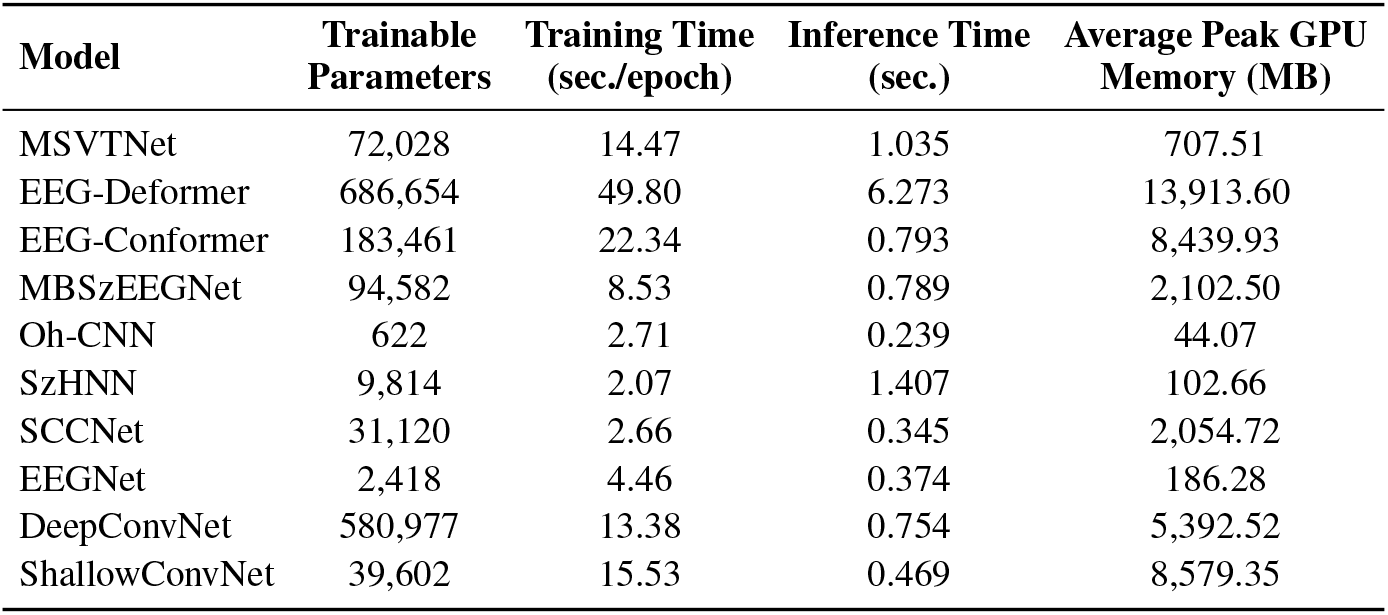
Computational efficiency of the evaluated models for the MCI vs. Normal classification task.

| Model | Trainable Parameters | Training Time (sec/epoch) | Inference Time (sec.) | Average Peak GPU Memory (MB) |
| --- | --- | --- | --- | --- |
| MSVTNet | 72,028 | 14.47 | 1.035 | 707.51 |
| EEG-Deformer | 686,654 | 49.80 | 6.273 | 13,913.60 |
| EEG-Conformer | 183,461 | 22.34 | 0.793 | 8,439.93 |
| MBSzEEGNet | 94,582 | 8.53 | 0.789 | 2,102.50 |
| Oh-CNN | 622 | 2.71 | 0.239 | 44.07 |
| SzHNN | 9,814 | 2.07 | 1.407 | 102.66 |
| SCCNet | 31,120 | 2.66 | 0.345 | 2,054.72 |
| EEGNet | 2,418 | 4.46 | 0.374 | 186.28 |
| DeepConvNet | 580,977 | 13.38 | 0.754 | 5,392.52 |
| ShallowConvNet | 39,602 | 15.53 | 0.469 | 8,579.35 |

**Table 18:**
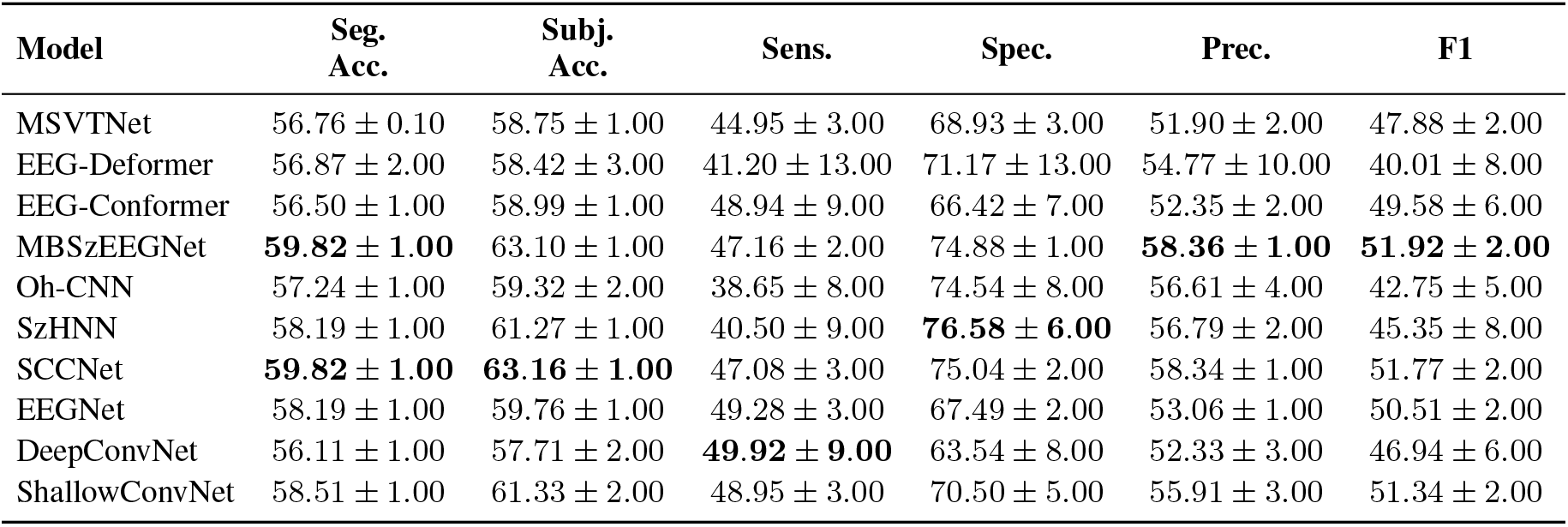
Performance comparison of deep learning models on the MCI vs. Dementia classification task. Values are reported as mean ± standard deviation in percentage points.

**Table 19:** Computational efficiency of the evaluated models for the MCI vs. Dementia classification task.

| Model | Trainable Parameters | Training Time (sec/epoch) | Inference Time (sec.) | Average Peak GPU Memory (MB) |
| --- | --- | --- | --- | --- |
| MSVTNet | 72,028 | 18.44 | 0.9328 | 708.35 |
| EEG-Deformer | 686,654 | 42.08 | 5.9619 | 13,913.60 |
| EEG-Conformer | 183,461 | 16.31 | 0.7076 | 8,440.55 |
| MBSzEEGNet | 94,582 | 7.71 | 0.7026 | 2,102.50 |
| Oh-CNN | 622 | 4.04 | 0.2082 | 44.07 |
| SzHNN | 9,814 | **1.87** | 0.8560 | 103.41 |
| SCCNet | 31,120 | 3.59 | 0.2863 | 2,054.72 |
| EEGNet | 2,418 | 4.04 | 0.3302 | 186.28 |
| DeepConvNet | 580,977 | 14.64 | 0.6739 | 5,392.92 |
| ShallowConvNet | 39,602 | 13.41 | 0.4155 | 8,579.35 |

